# Homozygous c.820G>A variant in MGME1 contributes to multi-systemic mitochondrial dysfunction in an Indian patient cohort

**DOI:** 10.64898/2026.05.18.725852

**Authors:** Pranavi Hegde, Amoolya Kandettu, Arpan Bhattacharyya, Budheswar Dehury, Sruthi S Nair, Rajalakshmi Poyuran, Smily Sharma, Benu Brata Das, Soumya Sundaram, Sanjiban Chakrabarty

**Affiliations:** Department of Public Health Genomics, Manipal School of Life Sciences, Manipal Academy of Higher Education, Manipal, Karnataka, India; Laboratory of Molecular Biology, School of Biological Sciences, Indian Association for the Cultivation of Science, 2A&B, Raja S. C. Mullick Road, Jadavpur, Kolkata 700032, India; Department of Bioinformatics, Manipal School of Life Sciences, Manipal Academy of Higher Education, Manipal, Karnataka, India; Department of Neurology, Sree Chitra Tirunal Institute for Medical Sciences and Technology, Trivandrum, Kerala, India; Department of Pathology, Sree Chitra Tirunal Institute for Medical Sciences and Technology, Trivandrum, Kerala, India; Department of Imaging Sciences and Interventional Radiology, Sree Chitra Tirunal Institute for Medical Sciences and Technology, Trivandrum, Kerala, India

**Keywords:** Mitochondrial disorder, MGME1, mtDNA replication, mtDNA maintenance disorder

## Abstract

Mitochondrial DNA (mtDNA) maintenance disorders arise from defects in mtDNA replication or repair, frequently resulting in extensive deletions or depletion of mtDNA. Mitochondrial genome maintenance exonuclease 1 (MGME1) is a nuclear-encoded nuclease essential for mtDNA replication and genome stability, and biallelic pathogenic variants in MGME1 cause mitochondrial DNA depletion syndrome 11. Here, we report a novel homozygous MGME1 missense variant c.820G>A (p. Ala274Thr) in five affected individuals from unrelated South Indian families presenting with proximal myopathy, chronic progressive external ophthalmoplegia, and cardiac and renal involvement. Patient-derived cells exhibited a significant reduction in mtDNA copy number, consistent with impaired mtDNA maintenance. Mechanistic analyses combining imaging-based and biochemical approaches demonstrated that the MGME1 variant disrupts both mtDNA replication and repair. Functional characterization further revealed defective oxidative phosphorylation and reduced mitochondrial membrane potential, confirming mitochondrial dysfunction. Collectively, our findings establish the pathogenicity of this novel MGME1 variant and expand the clinical and molecular spectrum of MGME1-associated mitochondrial disease, linking impaired mtDNA replication to multisystemic mitochondrial dysfunction.

**Graphical abstract summary:** Schematic overview of mitochondrial DNA (mtDNA) replication in wild-type and MGME1 mutant conditions. In wild-type cells, the coordinated activity of the mitochondrial replication machinery (TWNK, POLG, LIG3, TFAM, and MGME1) maintains mtDNA integrity and supports normal oxidative phosphorylation and ATP production. In contrast, the MGME1 mutation disrupts mtDNA replication and processing, leading to replication defects, mtDNA depletion, and mitochondrial dysfunction.

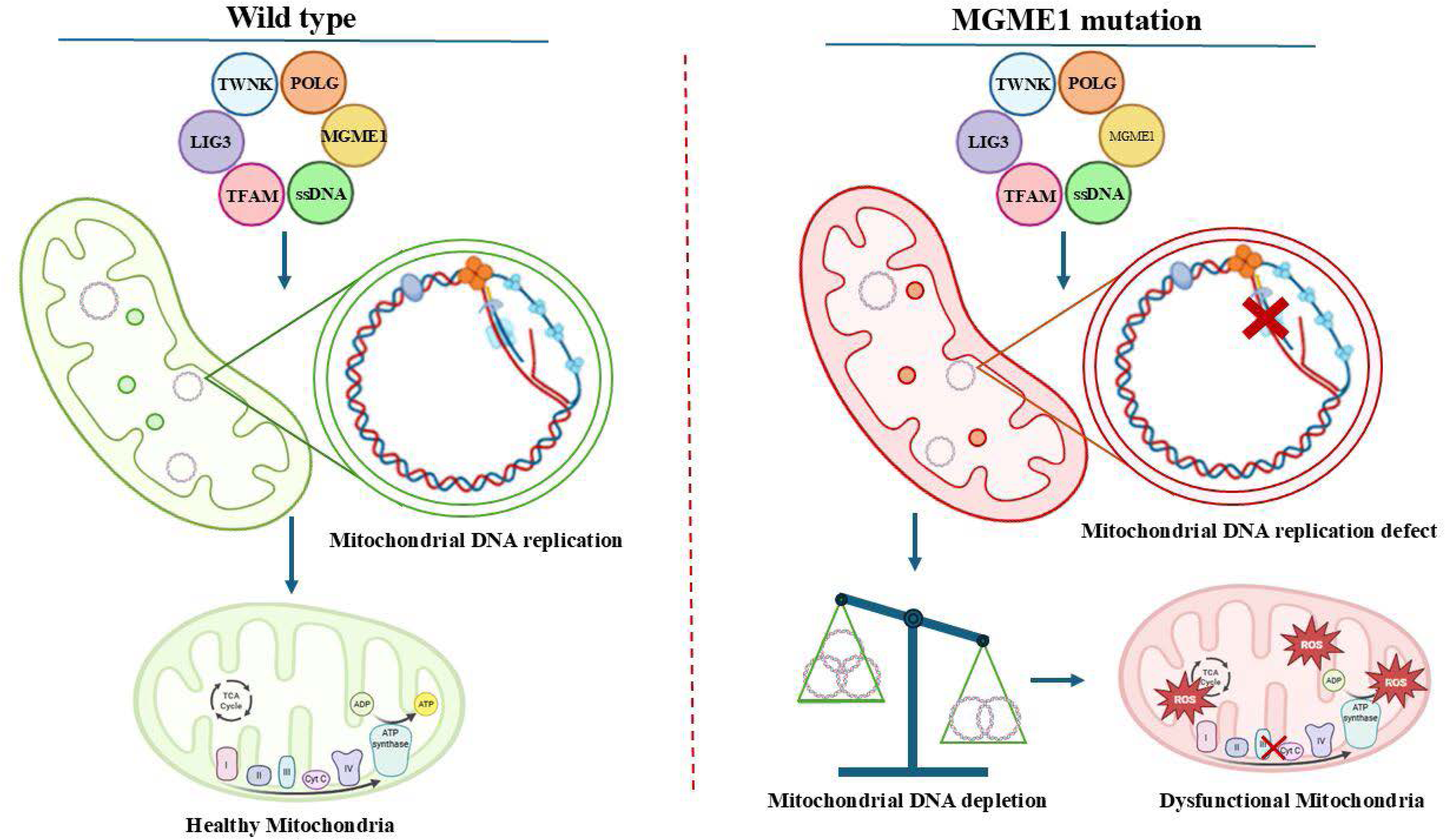

## Introduction

Mitochondrial DNA (mtDNA) is a circular genome of about 16.6 kb encoding 13 oxidative phosphorylation subunits. Maintenance, replication, and repair of mtDNA rely entirely on nuclear-encoded cytoplasmic proteins imported into mitochondria(Bhattacharjee & Das, 2025; Roy *et al*, 2022; Russell & Turnbull, 2014; Yan *et al*, 2019). Mitochondria lack comprehensive DNA replication and repair systems, relying on specialized pathways for mtDNA integrity, linked to mitochondrial dynamics, ensuring genome stability and cellular energy homeostasis(Akbari *et al*, 2022; Bhattacharjee & Das, 2025; El-Hattab & Scaglia, 2013). Nuclear-encoded proteins preserve mitochondrial genome integrity via the mitochondrial nucleotide salvage pathway, converting deoxyribonucleosides (dT, dC, dG, dA) into dNMPs via thymidine kinase 2 (TK2) and deoxyguanosine kinase (DGUOK), then phosphorylating to dNDPs by nucleotide monophosphate kinase (NMPK) and to dNTPs by nucleotide diphosphate kinase (NDPK). NDPK associates with succinyl-CoA ligase and γ-aminobutyrate transaminase, linking nucleotide homeostasis to mitochondrial metabolism. In the cytosol, ribonucleotide reductase (RNR), including the p53-inducible subunit RRM2B, generates dNDPs from ribonucleotides, while thymidine phosphorylase regulates thymidine turnover. Nucleotide transport into mitochondria is mediated by proteins like ANT1, AGK, and MPV17, supplying substrates for mtDNA synthesis(Almannai *et al*, 2022). Mitochondrial DNA replication and repair are executed by specialized nuclear-encoded machinery, including polymerase γ (POLG), TWINKLE helicase, mitochondrial single-stranded DNA-binding protein (mtSSB), and mitochondrial RNA polymerase (POLRMT). Genome integrity is safeguarded by nucleases such as APE1, APE2, DNA2, FEN1, EXOG, ENDO G, and MGME1, which process replication and repair intermediates to maintain mtDNA stability(Bruni *et al*, 2017). Mitochondrial Genome Maintenance Exonuclease 1 (MGME1) is a mitochondrial DNA nuclease in the Rec B family. This protein processes 5′ and 3′ flaps at mtDNA replication ends, essential for removing RNA primers at replication origins (OriH and OriL) (Gonzalez *et al*, 2024; Mao *et al*, 2024). MGME1 binds single-stranded DNA, efficiently cleaving long 5′ flaps, contributing to replication intermediate maturation and preventing unligatable nicks. Mgme1−/− mice show reduced mtDNA copy number and deletion accumulation; despite significant mitochondrial dysfunction, they remain viable and do not develop progeroid features, suggesting MGME1-associated defects are distinct from aging-related mtDNA instability(Kornblum *et al*, 2013; Milenkovic *et al*, 2022; Matic *et al*, 2018). Although MGME1 is implicated in processing replication intermediates to preserve mtDNA integrity, its precise role in human disease remains incompletely understood(Uhler *et al*, 2016). Here, we describe an Indian cohort with five unrelated individuals harboring a novel homozygous MGME1 missense variant c.820G>A (p. Ala274Thr) with multisystem mitochondrial disease.

## Results

### Clinical presentation of subjects with the MGME1 mutation

#### Patient 1

Patient 1, a 44-year-old female, had experienced bilateral ptosis and proximal muscle weakness for 4 years. Over the last two years, she had four episodes of respiratory failure and worsening muscle weakness, leading to invasive ventilation, with two of these incidents triggered by pneumonia. The patient was initially diagnosed with myasthenia gravis and subsequently administered corticosteroids, intravenous immunoglobulin, plasma exchange, and pyridostigmine, which resulted in partial recovery. The patient had a history of diabetes mellitus, hypertension, and asthma, which predated the onset of neurological symptoms. A recent examination revealed myopathic facies, bilateral fatigable ptosis, restricted eye movements, and bifacial weakness. Muscle strength was assessed as follows: neck muscles (grade 3); shoulder flexion and abduction (grade 4+); hip flexion (grade 4−); hip adduction and abduction (grade 4+); and knee flexion (grade 4+). Tone, sensory, and cerebellar examinations were normal. Her previous photographs revealed the presence of ptosis from the age of 30, and she had an unremarkable birth and developmental history. The results of her routine hemogram, renal function, and liver function tests were all normal. Additionally, her serum creatinine kinase (CK) and lactate levels were within the normal ranges. However, her thyroid-stimulating hormone (TSH) level was elevated to 10.02 µIU/ml. Her workup for myasthenia gravis, including anti-acetylcholine receptor and muscle-specific tyrosine kinase antibodies, as well as repetitive nerve stimulation (RNS) tests of the facial, spinal accessory, and ulnar nerves, was negative. Nerve conduction study (NCS) and electromyography (EMG) results were normal (Table 1). MRI (Magnetic resonance imaging) of the brain showed no abnormalities (Supplementary Fig. 1) while of the muscles revealed fatty replacement of the vastus intermedius and patchy involvement of the glutei, hamstrings, soleus, and erector spinae muscles (Supplementary Fig. 2). Muscle biopsy revealed preserved fascicular architecture with myopathic features (Fig. 1A). Modified Gomori Trichrome (MGT stain) revealed ragged red fibres, and many COX-deficient fibres were highlighted in the COX-SDH staining (Fig. 1B and C). NADH-TRs occasionally appeared as ragged blue fibers, and Oil Red O staining revealed the presence of neutral lipid accumulation in occasional fibers (Fig. 1D and E). Type 1 fiber predominance was observed by ATPase staining at pH 9.5. Electron microscopy of the muscle tissue revealed abnormal mitochondria (Fig. 1F).

**Table 1:**
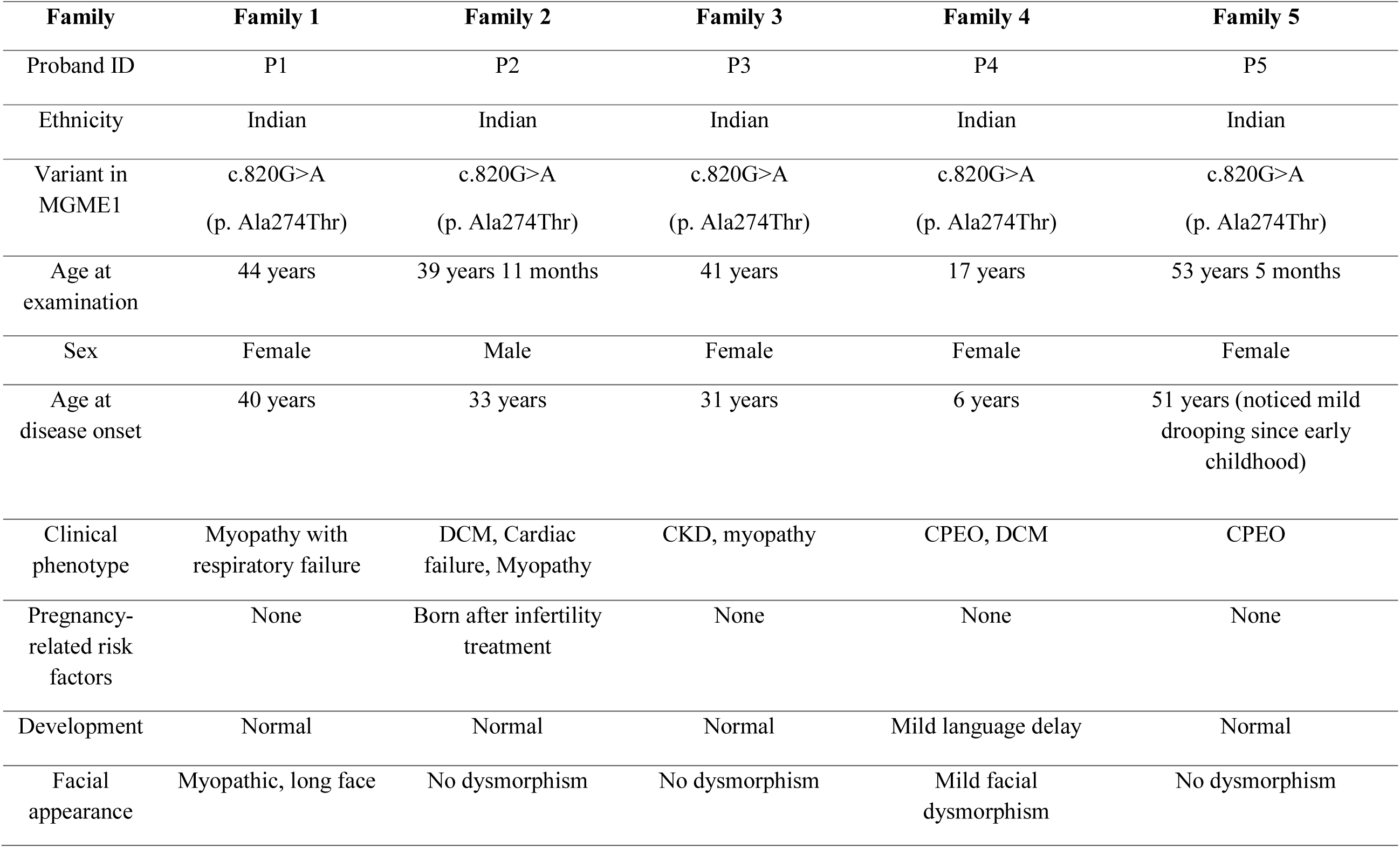
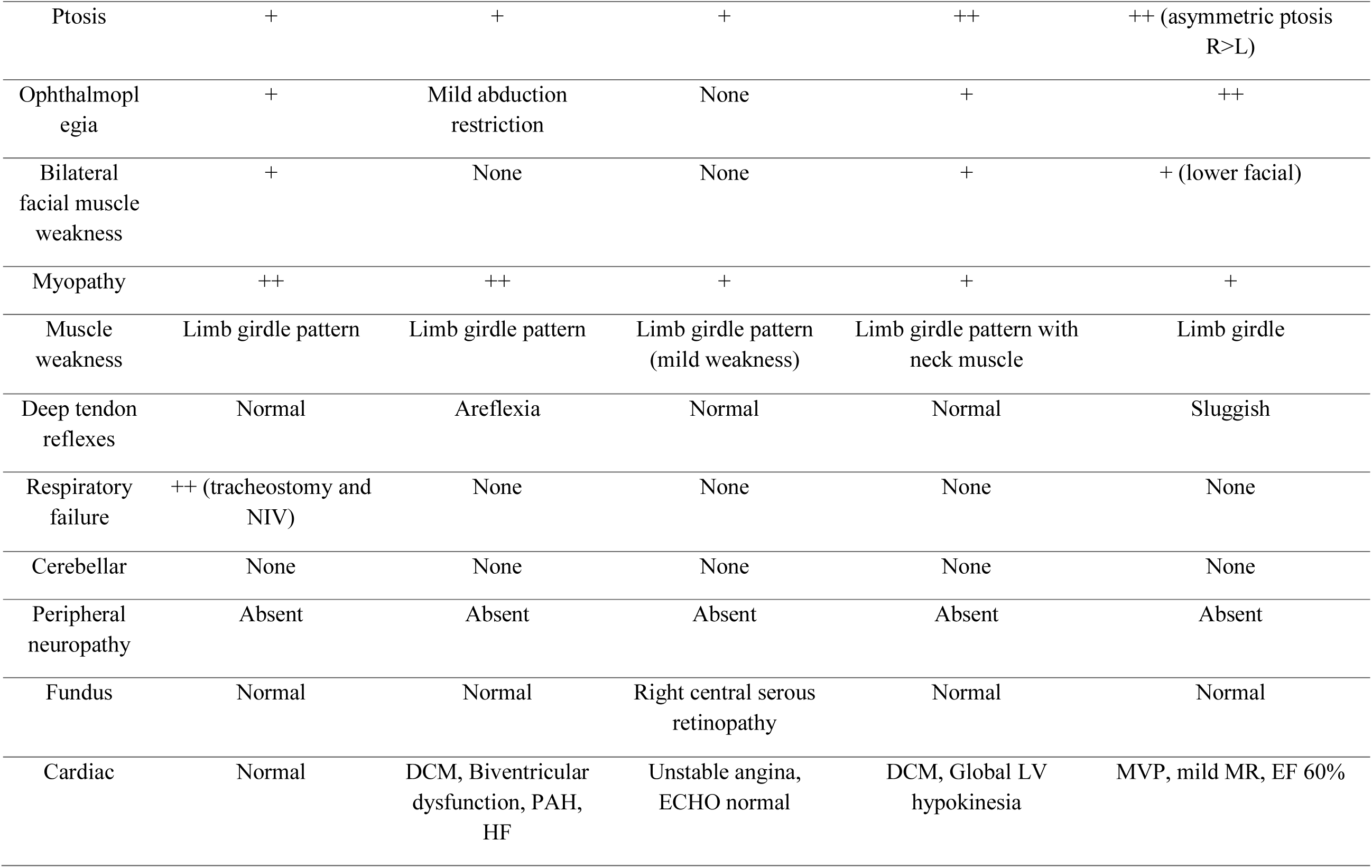
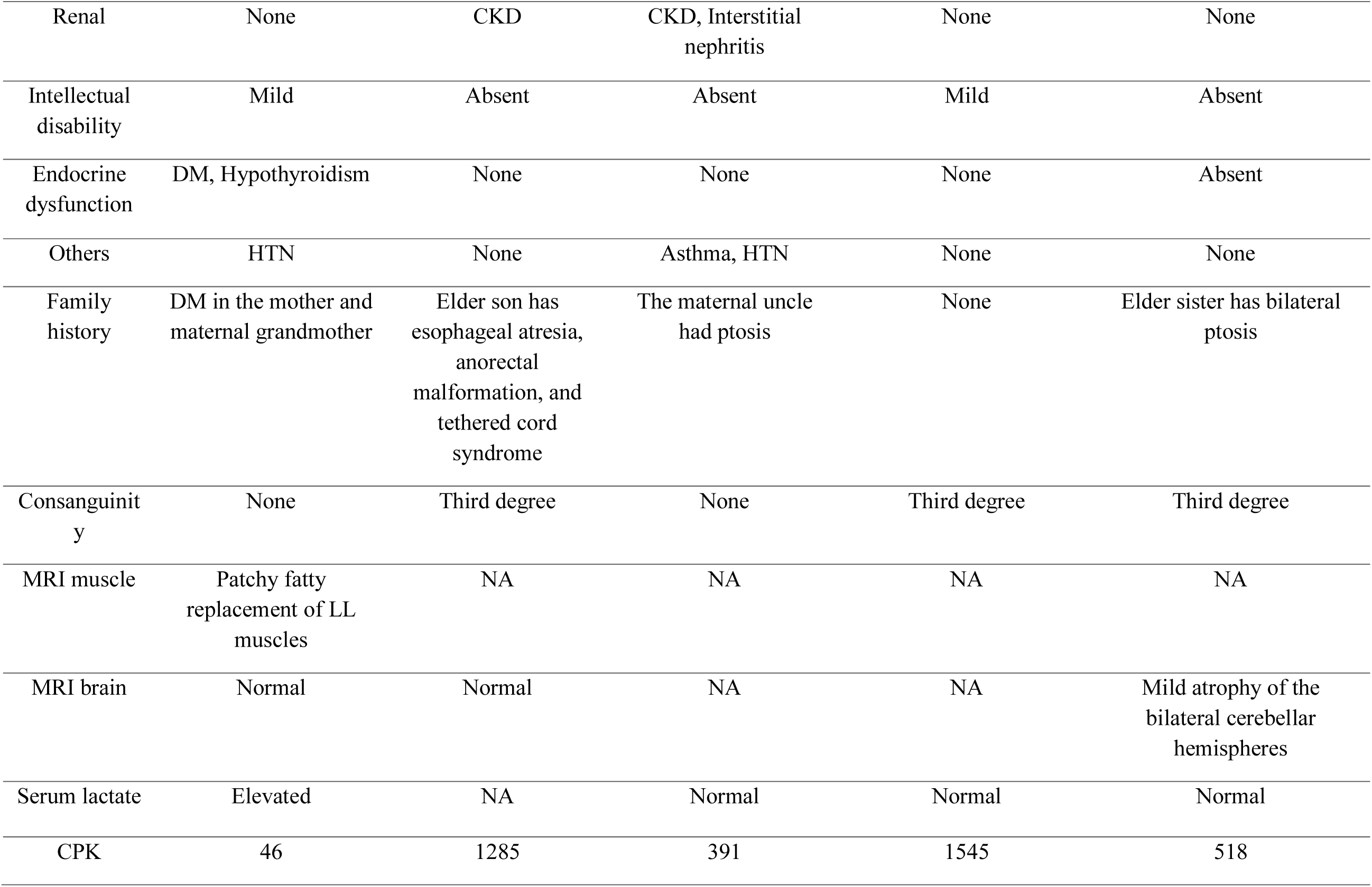
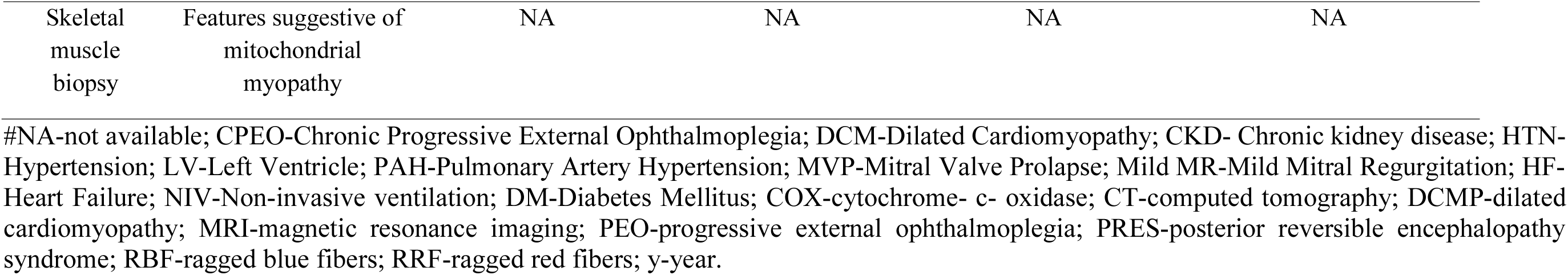
Clinical spectrum of patients with MGME1 variation in the present study.

**Figure 1.**
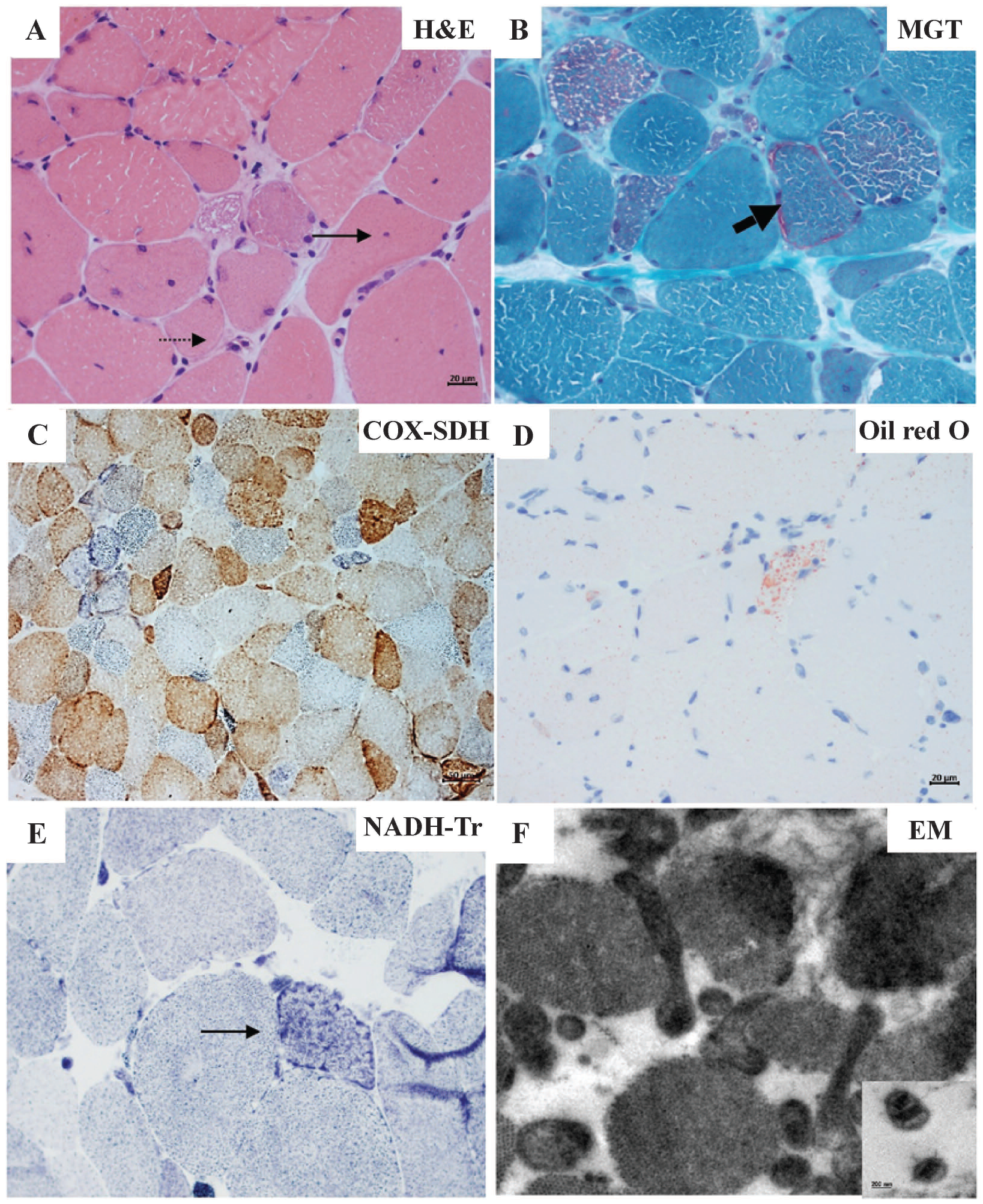
Mitochondrial enzyme histochemistry and electron microscopy analysis of the muscle biopsy of patient 1 with the MGME1 variant. A) Hematoxylin and Eosin (H & E) stain showing rounding of fibres, variation in fibre size, atrophic fibres and fibres with internalized nuclei (arrow), scale bar 20 µm; B) Modified Gomori Trichrome (MGT) stain showing ragged red fibres (thick arrow), scale bar 20 µm; C) Many COX-deficient fibres (blue fibres) in COX-SDH stain, scale bar 50 µm; D) Muscle fibre with fine lipid-filled vacuoles in Oil Red O, scale bar 20 µm; E) Ragged blue fibre in Nicotinamide Adenine Dinucleotide Tetrazolium Reductase (NADH-Tr) stain; F) Electron microscopy showing abnormal shape of mitochondria and paracrystalline inclusions.

#### Patient 2

Patient 2, a 39-year-old male, presented with dyspnoea on exertion, orthopnoea, chest pain, and palpitations for 6 years. The patient was diagnosed with dilated cardiomyopathy and heart failure. He subsequently had difficulty climbing stairs and rising from a squatting position, suggesting proximal lower limb muscle weakness without myalgia or muscle cramps. He is the only child of third-degree consanguineous parents and achieves developmental milestones at an appropriate age. He had mild drooping of both eyelids since early childhood. His older son had oesophageal atresia, anorectal malformation, congenital talipes equinovarus deformity, ventricular septal defects with spontaneous closure, and tethered cord syndrome. Upon examination, bilateral partial ptosis and mild restriction of lateral eye movement were noted. He had mild weakness in the elbow extensors, hip adductors, abductors, flexors, and extensors. Mild wasting of the bilateral thenar, hypothenar, deltoid, pectoralis, and forearm muscles was observed. The patient had hypotonia and areflexia. His hemogram and liver and thyroid function tests were normal. Renal function tests revealed evidence of chronic kidney disease (blood urea nitrogen, 43 mg/dL; serum creatinine, 2.05 mg/dL). His serum CPK level was elevated, and electromyography (EMG) showed myopathic potential. The patient was negative for NCS and RNS. Echocardiography revealed dilated cardiomyopathy (DCM) with poor ejection fraction. Cardiac MRI confirmed DCM with moderate biventricular dysfunction, global subendocardial late gadolinium enhancement of the myocardium with an increased ECV fraction, pulmonary arterial hypertension, and mild pericardial and pleural effusions (Table 1) (Supplementary Fig. 3).

#### Patient 3

Patient 3 was a 41-year-old female who presented with a history of 10-year right-eyelid ptosis and intermittent diplopia. The patient did not experience any improvement after pyridostigmine treatment. Her medical history revealed chronic kidney disease due to interstitial nephritis and unstable angina, which was diagnosed one year prior to presentation. Additionally, her maternal uncle had ptosis, and three neonatal and childhood deaths occurred in her paternal family. Upon examination, bilateral ptosis and mild weakness of the proximal upper and lower limb muscles (grade 4+) were observed with normal tone and deep tendon reflexes. Sensory and cerebellar examination results were normal. Laboratory tests revealed a serum creatinine level of 1.71 mg/dL and a CK level of 391 U/L, confirming mild elevations. Routine hemogram and liver and thyroid function tests were normal. Abdominal ultrasonography revealed bilateral Grade II renal disease, and NCS and RNS were normal. Fundus examination revealed right central serous retinopathy. Cardiac evaluations, including an electrocardiogram, Holter monitoring, and echocardiogram, were normal (Table 1).

#### Patient 4

Patient 4 was a 15-year-old girl born to third-degree consanguineous parents. She had a normal birth but a mild delay in language milestones. Since the age of 6 years, she has noted bilateral ptosis. Upon evaluation, the patient had a nasal twang in her speech, bilateral complete ptosis, severe restriction of eye movements, bifacial weakness, and an absent gag reflex. She had weakness in her neck muscles (grade 4), shoulder flexors (grade 4), abductors (grade 4+), hip flexors (grade 4-), adductors (grade 4), extensors (grade 4), abductors (grade 4+), and dorsiflexors of the ankle (grade 4+). Her hemogram and routine blood biochemistry results were normal, but her CK level was elevated (1545 U/L). Serum lactate levels were normal (1.7 mg/dl). The NCS was normal, and the EMG showed myopathic potential. Initially, cardiac evaluation, including echocardiogram and Holter monitoring, revealed no abnormalities. However, two years later, she developed exertional dyspnoea and was diagnosed with dilated cardiomyopathy. Echocardiography revealed global hypokinesia, an ejection fraction of 18%, and mild mitral and tricuspid regurgitation (Table 1).

#### Patient 5

Patient 5 was a 53-year-old woman, born to third-degree consanguineous parents, who presented with long-standing asymmetrical bilateral ptosis (right greater than left) of 10–15 years duration, without diplopia or fatigability. A positive family history was noted, as her elder sister also had bilateral ptosis. Neurological examination revealed severe restriction of extraocular movements bilaterally, along with ptosis, mild weakness of neck flexion, and proximal weakness of both upper and lower limbs. Routine haematological and biochemical investigations were within normal limits. Serum lactate was normal (1.1 mmol/L), while serum CK was mildly elevated (518 U/L). NCS, EMG, and RNS were normal. Cardiac evaluation did not reveal any abnormalities. Brain MRI of patient 5 demonstrated bilateral symmetrical cerebellar hemispheric atrophy, without a lactate peak on magnetic resonance spectroscopy (Table 1).

### Identification of the *MGME1* variant c.820G>A in patient samples

Singleton exome sequencing of probands P1, P2, P3, P4, and P5 revealed a novel homozygous missense variant, c.820G>A (p. Ala274Thr), in exon 4 of *MGME1* (NM_016034.5) (Fig. 2A). *In silico* analysis using MutationTaster2(Schwarz *et al*, 2014) and ClinPred(Alirezaie *et al*, 2018) consistently predicted that these variants impaired protein function. According to the American College of Medical Genetics and Genomics(Richards *et al*, 2015), the c.820G>A (p. Ala274Thr) variant was classified as a variant of uncertain significance.

**Figure 2.**
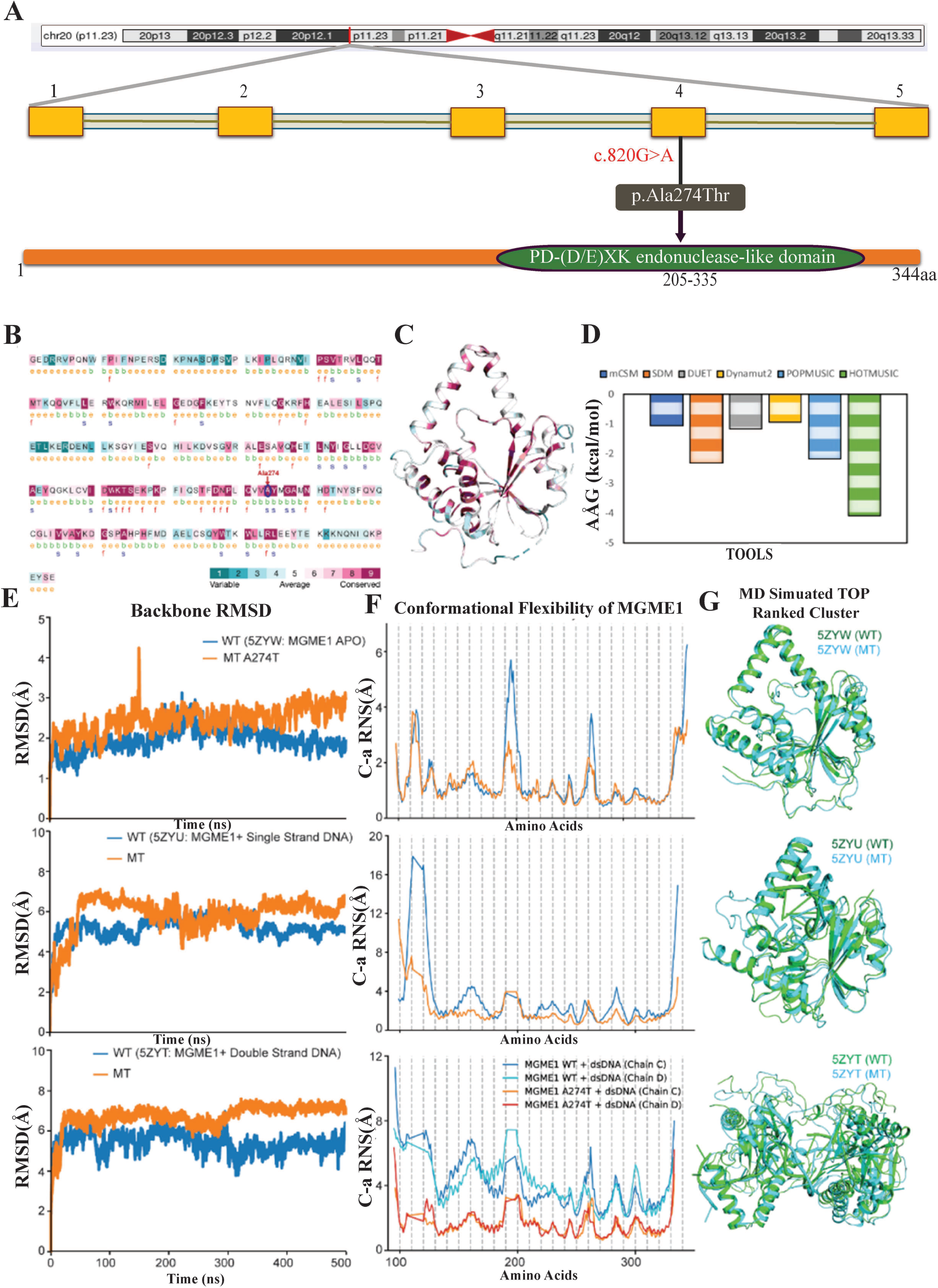
Sequence-structure conservation analysis and effect of variation (A274T) on protein stability and its conformational dynamics in MGME1. A) Variation mapped on the domains of MGME1. Variants are indicated in red (https://pfam.xfam.org/); B) Conservation analysis of MGME1 at sequence level displaying variable, average and conserved residues; C) CONSURF analysis displaying the conserved (purple) and variable (cyan) residues in apo structure of MGME1 (Ala274 a strongly conserved residue is shown in stick format); D) Assessing the effect of variation on change in protein stability (ΔΔG) measured by various structure-based protein stability tools like mCSM, SDM, Dynamut2, POPMUSIC and HOTMUSIC; E) Backbone root mean square deviation (RMSD) of the wild type and variant A274T of MGME1 in apo (PDB ID: 5ZYW), single strand DNA bound (5ZYU) and with 3’ overhang double strand DNA bound states (5ZYT) over a time scale of 500 ns assessed using all-atoms molecular dynamic simulations; F) Conformational flexibility of the wild type and variant states of MGME1 displaying the root mean square fluctuations of Cα atoms measured from the last 300 ns of MD trajectories; G) Structural overlay of the top ranked cluster representatives of the wild type and variant type structures extracted from MD trajectories of MGME1 systems displaying structural changes in apo, single strand and double strand DNA bound.

### Conservation and protein stability predictions of the MGME1 variant

Sequence conservation analysis revealed that Ala274 lies within a strongly conserved stretch of MGME1, positioned in the α-helical region that forms a part of the protein’s core structural framework (Fig. 2B). Mapping conservation scores onto the three-dimensional structure confirmed the importance of the residue within a conserved patch within MGME1 (Fig. 2C). Sequence analysis of related family members further highlighted the conserved secondary structural features across species, positioning Ala274 within helix α5 as a structurally sensitive site (Supplementary Fig. 4). Previous studies have established MGME1 as an essential regulator of mitochondrial DNA maintenance, with variation in conserved residues leading to impaired enzymatic activity and genome instability(Yang *et al*, 2018).

Multiple protein stability prediction tools were used to assess the effect of A274T substitution which consistently indicated destabilization, with negative ΔΔG values obtained from mCSM (−1.05 kcal/mol), SDM (−2.32 kcal/mol), DUET (−1.17 kcal/mol), and Dynamut2 (−0.94 kcal/mol), confirming the destabilizing effect (Fig. 2D). Although the normal mode analysis yielded a slightly positive ΔΔG_ENCoM (0.219 kcal/mol), vibrational entropy analysis indicated a reduction in molecular flexibility (ΔΔS_vib = −0.274 kcal/mol/K). These results underscore the functional significance of Ala274 and align with recent structural findings showing that MGME1 adopts a rigid DNA-clamp conformation when bound to 5′ overhang DNA, a configuration that depends on conserved residues to maintain stability and function(Mao *et al*, 2024).

### Conformational dynamics of MGME1 variant

To explore the structural consequences of the variation, 500 ns molecular dynamics simulations were performed for wild-type and A274T MGME1 in the apo, ssDNA-bound, and dsDNA-bound states. The backbone RMSD values from the MD simulations showed that the wild-type protein occupied a broader conformational space, whereas the variant protein was restricted to narrower conformations, reflecting reduced flexibility (Fig. 2E). The RMSF profiles, calculated from the last 300 ns of the MD trajectories, supported this finding, with the variant consistently displaying dampened residue-level fluctuations compared with those of the wild type (Fig. 2F).

Compared with the wild-type complex, the variant formed a greater number of hydrogen bonding interactions with both single-stranded DNA (ssDNA) and double-stranded DNA (dsDNA). However, these additional bonds showed altered occupancy with a high degree of instability, indicating a more rigid, yet less adaptable binding mode (Supplementary Fig. 5). Structural clustering of MD-simulated snapshots further highlighted distinct conformational ensembles, and overlays of representative top-ranked clusters revealed rearrangements in DNA-binding loops and local secondary structure elements in the variant compared with the wild type (Fig. 2G). These findings indicate that the A274T variant reduces conformational adaptability and introduces rigidification within the DNA-binding regions. This rigidity may impair DNA processing, consistent with observations in other mitochondrial nucleases, such as EXOG, where the flexibility of the DNA-binding domains is essential for engaging flap structures and ensuring efficient processing(Karlowicz *et al*, 2025).

### MGME1 variant contributes to decreased MGME1 expression in patient-derived cells

Quantitative real-time PCR demonstrated reduced *MGME1* expression in both P1 and P2 compared to Control cells (Fig. 3A). Immunoblot analysis revealed a lower abundance of MGME1 protein in P1 and P2 cells than in the Control cells (Fig. 3B). Analysis of mtDNA copy number by qRT-PCR confirmed mtDNA depletion in P1 and P2 patient cell lines (Fig. 3C).

**Figure 3.**
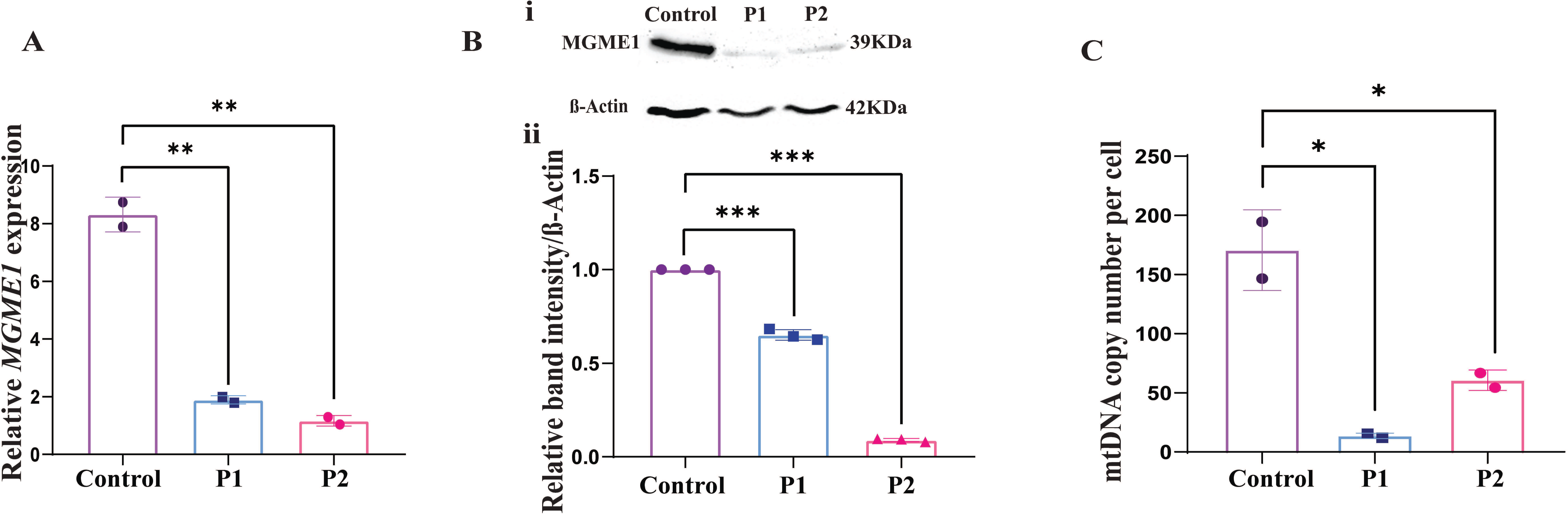
MGME1 variant contributes to decreased MGME1 expression and mtDNA copy number. A) qRT-PCR analysis of the *MGME1* in Control, P1, and P2 cells; β-Actin was used as an endogenous control. Relative quantity (RQ) was calculated using the formula 2^−ΔΔCT^, **p < 0.01(n=3); B) (i)Western blot analysis of MGME1 protein in Control, P1, and P2. Representative blot images of MGME1 protein and β-Actin; (ii) Densitometric analysis was performed after normalizing the MGME1 protein band intensity to the respective β-Actin, ***p < 0.001 (n=3); C) Bar graph showing reduced relative mitochondrial DNA copy number in P1 and P2 compared to control cells, **p<0.01(n=3). Data are presented as mean ± SEM (Student’s t-test).

### Loss of MGME1 expression contributes to decreased mtDNA copy number and nucleoid

*In silico* prediction from molecular dynamics simulations indicated that the A274T mutation destabilises MGME1 by introducing rigidity in the DNA-binding region (Supplementary Fig. 5). To gain a more in-depth understanding of mtDNA stability in p. Ala274Thr patients, we performed super-resolution microscopy of patient fibroblasts to examine mitochondrial length and nucleoid number. Mitochondrial DNA depletion syndrome is characterised by stressed mitochondria that bear with compromised energy metabolism under disease states. These dysfunctional mitochondria exhibit altered mitochondrial morphology, ranging from fragmented or fissioned to hyperfused. P1 patient fibroblasts represented severely hyperfused mitochondria as compared to Control, with mitochondrial lengths several-fold elevated (p < 0.0001) in patient cell lines in log scale (Fig. 4Ai). Furthermore, we observed a significant reduction in mitochondrial nucleoids per cell in P1 patient fibroblasts compared to control, indicative of severe mtDNA depletion (Fig. 4Aii). Consequently, these patient cells exhibit mtDNA depletion syndromes, characterized by hyperfused mitochondria with sparsely distributed nucleoids, indicative of stressed mitochondria. We conducted immunofluorescence analysis to evaluate the expression of TFAM, a mitochondrial nucleoid protein, and observed reduced TFAM levels in P1 cells. These findings imply that MGME1 variation disrupts mtDNA replication and affects nucleoid numbers (Fig. 4B).

**Figure 4.**
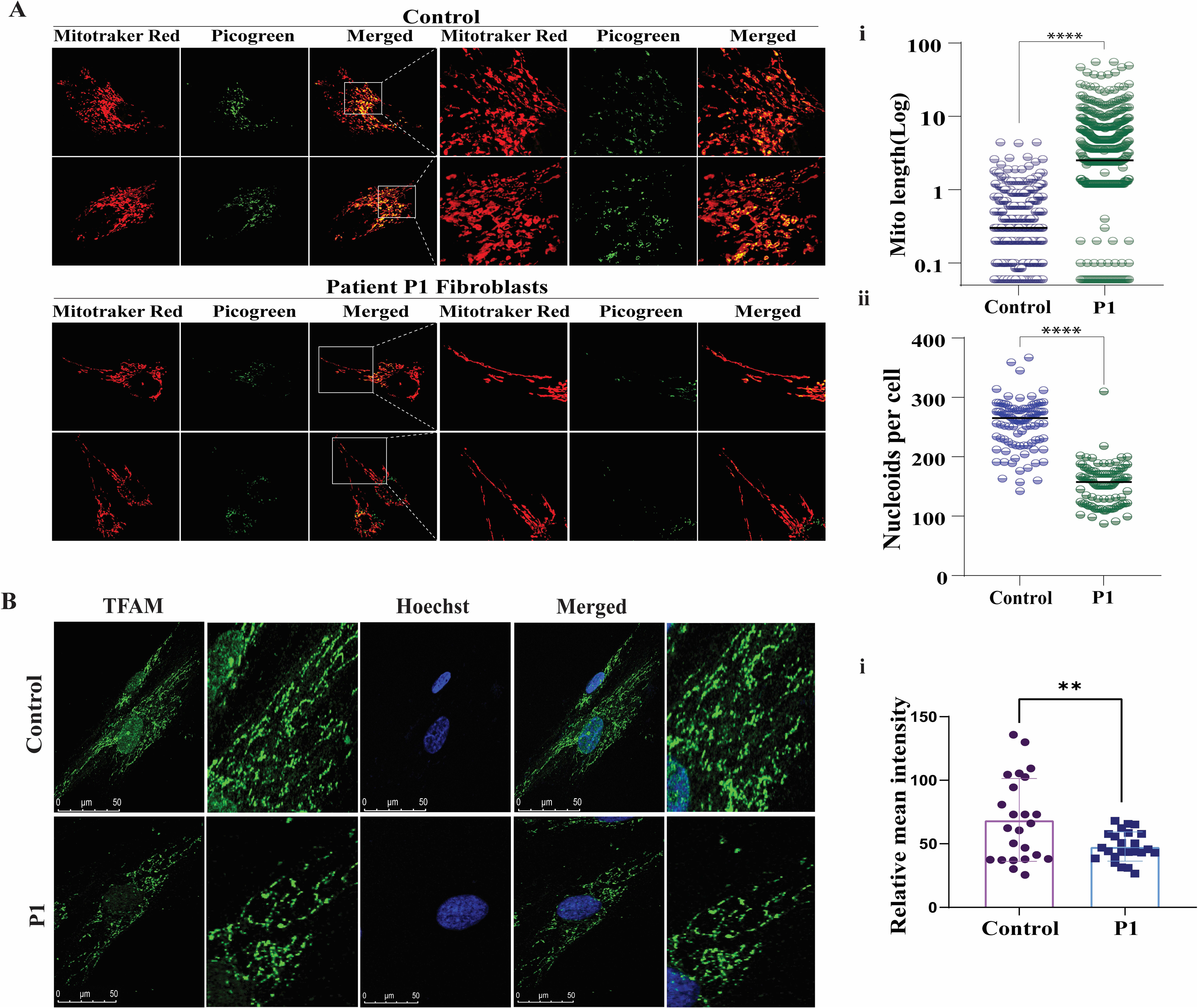
MGME1 variant contributes to decreased nucleoid content. A) Representative confocal image of Control and P1 stained with PicoGreen and MitoTracker Red, showing i) elongated mitochondria with ii) reduced mitochondrial nucleoids per cell, ****p<0.0001(n=3); B) Representative confocal images of immunofluorescence analysis of TFAM protein; i) Bar graph showing reduced relative intensity of TFAM protein in MGME1 variant patient-derived P1 cells, **p<0.01(n=3). Data are presented as mean ± SEM (Student’s t-test).

### MGME1 variant contributes to impaired mtDNA replication and repair

We investigated the mtDNA replication capacity using 2′,3′-dideoxycytidine (ddC) in control and patient derived P1 cells. Upon drug withdrawal, the control cells exhibited a marked recovery of mtDNA, whereas no such recovery was observed in P1 cells, indicating a decreased mtDNA replication capacity in P1 cells (Fig. 5A). Subsequently, the mtDNA repair capacity was analysed in control and P1 fibroblasts. Control cells showed reduced PCR amplification immediately after H₂O₂ treatment, followed by marked recovery after 6 h, indicating efficient repair of mtDNA damage. In contrast, P1 cells exhibited minimal recovery of intact mtDNA, suggesting severe impairment in mtDNA repair capacity (Fig. 5B-C).

**Figure 5.**
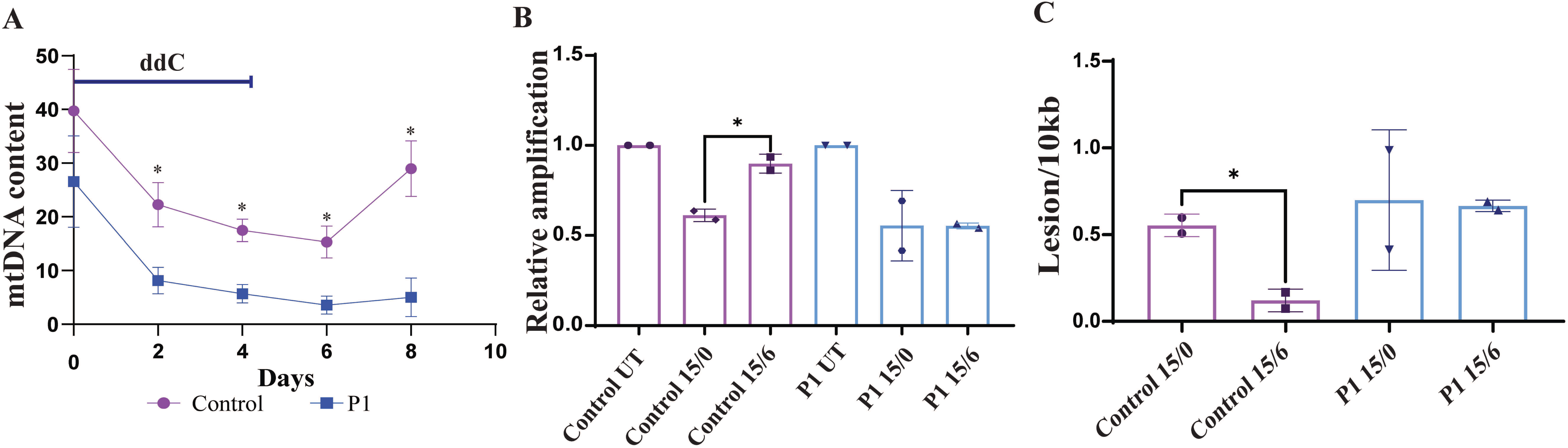
MGME1 variant impairs mitochondrial DNA replication and repair capacity. A) Mitochondrial DNA (mtDNA) depletion assay. Control and P1 cells were treated with ddC for 6 days. Relative mtDNA content was quantified at the indicated time points (2, 4, 6, 8 days). Control cells exhibit a progressive decrease in mtDNA copy number during ddC treatment, followed by partial recovery, whereas patient (P1) cells show a more pronounced depletion and impaired recovery kinetics, indicating compromised mtDNA replication capacity, *p<0.05 (n=3); B) Assessment of mtDNA repair following oxidative stress. Control and P1 cells were either untreated (UT), exposed to H₂O₂ for 15 minutes without recovery (15/0), or exposed to H₂O₂ followed by a 6-hour recovery period (15/6). mtDNA damage and repair were assessed using a long-amplicon PCR assay. Untreated cells (UT) served as baseline controls. H₂O₂ treatment significantly reduced amplification in both groups at 15/0, reflecting oxidative lesion formation. After 6 hours of recovery, P1 cells demonstrate significantly reduced recovery compared to control, indicating impaired mtDNA repair kinetics, *p < 0.05 (n=3); C) Quantification of mtDNA lesion frequency. The number of mtDNA lesions per 10 kb was calculated based on long-amplicon PCR efficiency. Control cells showed a significant reduction in lesion frequency following recovery, whereas lesion burden remained elevated in P1 cells, confirming defective mitochondrial genome repair associated with the MGME1 variant, *p < 0.05 (n=3). Data are presented as mean ± SEM (Student’s t-test).

### MGME1 variant leads to mitochondrial dysfunction

Immunoblot analysis of the mitochondrial respiratory chain complex proteins demonstrated a significant decrease in the expression of complex I, III, and IV subunits in P1 and P2 cells compared to the control. In addition, a mild reduction in complex V expressions was observed in P1 cells, whereas P2 cells did not show any difference in complex V levels (Fig. 6Ai-iv). These observations were further validated by significant decreases in complex IV enzyme activity and decreased ATP levels in P1 and P2 cells indicating impaired mitochondrial OXPHOS and ATP synthesis (Supplementary Fig. 6). Cellular respiration analysis using Seahorse XF24 Extracellular Flux Analyzer confirmed a decreased OCR and ECAR, suggesting perturbed mitochondrial respiratory capacity in P1 cells with the MGME1 variant (Fig. 6B and C, Supplementary Fig. 7). Assessment of MMP using TMRM showed a marked reduction in MMP in P1 cells, reflecting impaired mitochondrial bioenergetic function (Fig. 6D). In parallel, mitochondrial ROS levels were significantly decreased (Fig. 6E).

**Figure 6.**
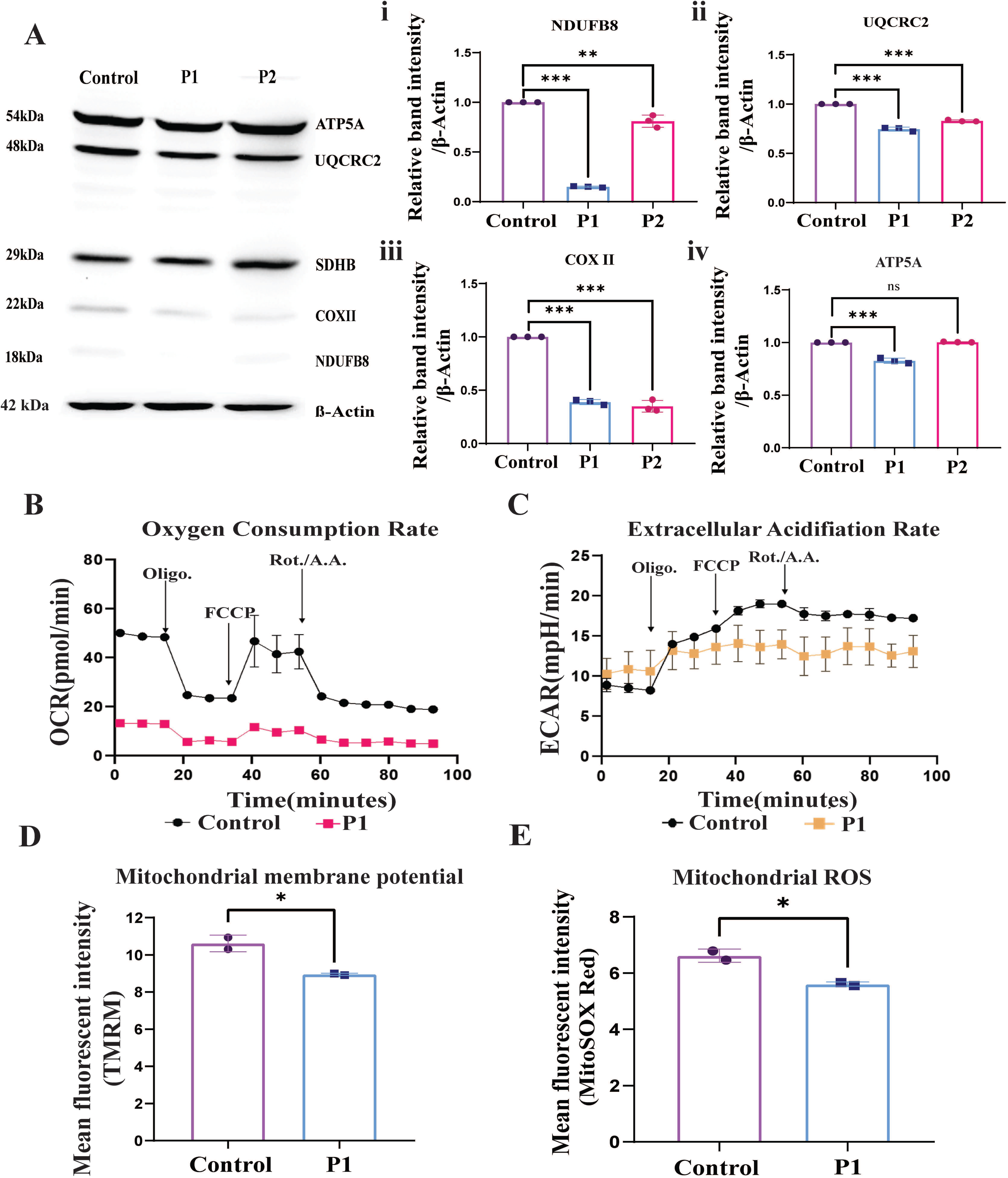
Mitochondrial dysfunction in patient-derived cells due to the MGME1 variant. A) Immunoblot images for protein expression of OXPHOS proteins in control and patient fibroblast (P1 and P2). (i-iv) Densitometric analysis of protein expression of NDUFB8 (Complex I), UQCRC2 (Complex III), COX II (Complex IV) and ATP5A (Complex V) in Control, P1 and P2 cells(n=3); B) Representative graph showing oxygen consumption rate (OCR) in control and P1 cell lines with injection of oligomycin, FCCP, and antimycin A and rotenone(n=3); C) Representative graph showing the extracellular acidification rate (ECAR) in control and P1. with injection of oligomycin, FCCP, and antimycin A and rotenone(n=3); D) Bar graph showing the reduced relative fluorescence intensity of Tetramethylrhodamine, methyl ester (TMRM) indicating decrease in mitochondrial membrane potential in P1 cell lines, *p<0.05; E) Bar graph showing the reduced relative fluorescence intensity of MitoSOX red indicating decrease in mitochondrial reactive oxygen species in P1, *p<0.05(n=3). Data are presented as mean ± SEM (Student’s t-test).

## Discussion

MGME1 is a major exonuclease that is involved in the maintenance and stabilization of mtDNA replication. Pathogenic mutations affecting the function of MGME1 have been shown to cause mtDNA depletion syndrome leading to multisystemic mitochondrial disease. In this study, we identified homozygous c.820G>A variation in the MGME1 gene in five unrelated patients with variable clinical phenotypes. Except for one patient who had disease onset at 6 years of age, the remaining patients presented in the fourth or fifth decade, highlighting that MGME1 should be considered in the differential diagnosis of late-onset mitochondrial myopathies. Consistent with previously reported literature, ptosis, ophthalmoplegia, and myopathy were the most frequent presenting features in our cohort.

Beyond skeletal muscle involvement, cardiac manifestations, including cardiomyopathy and cardiac failure, have been reported in MGME1-related disease and in our series, underscoring the importance of systematic cardiac evaluation and longitudinal surveillance in affected individuals. Renal involvement, as observed in two of our patients, has also been reported, with some patients initially presenting with chronic kidney disease and later developing CPEO or myopathy. Recurrent respiratory failure represents another potentially life-threatening complication and may clinically mimic myasthenia gravis, particularly in patients presenting with ptosis and ophthalmoplegia, potentially leading to diagnostic confusion. Additional clinical features reported in association with MGME1 mutations include cataracts, emaciation, and scoliosis. From a neuroimaging perspective, cerebellar atrophy has been the most consistently reported abnormality; however, only one patient in our series showed mild cerebellar volume loss. The phenotypic diversity in terms of clinical features, age of onset, and also complications observed in our case series with the same genetic variant is striking, necessitating in-depth phenotyping and monitoring for multi-organ involvement.

MGME1 c.820G>A variant disrupts mitochondrial DNA replication and impairs mitochondrial maintenance in these patients. MGME1, a nuclease encoded by nuclear genes and localized to mitochondria, is essential for managing the 5′ and 3′ flaps produced during replication of mtDNA. This activity is essential for the efficient removal of RNA primers from the origin of replication. The identification of a novel missense variant, c.820G>A, in these five cases supports accumulating evidence that pathogenic variation in MGME1 adversely affects mtDNA replication and overall stability.

A notable observation was the clinical variability among individuals carrying the same MGME1 variant. Our patient cohort with MGME1 c.820G>A (p. Ala274Thr) missense variant showed considerable differences in age onset, disease severity, and the extent of systemic involvement, indicating that genotype alone is insufficient to predict the clinical outcome. This variability likely reflects the influence of additional modifying factors, including nuclear genetic background and mitochondrial haplotype. In contrast, truncating variants such as c.456G>A (p. Trp152*) and c.862C>T (p. Gln288*) are associated with earlier disease onset, neurodevelopmental impairment, cerebellar atrophy, and more severe multisystem involvement, consistent with a greater degree of functional loss of MGME1 (Supplementary Table 1).

Our results showed a marked reduction in mtDNA copy number in cells harbouring the MGME1 variant, consistent with earlier studies of mitochondrial depletion syndromes associated with the MGME1 variant. The depletion was likely due to affecting the processing of mtDNA replication, leading to the accumulation of nicks that cannot be ligated, resulting in truncated mtDNA fragments and disrupting mitochondrial replication accuracy.

We further investigated whether this mtDNA replication defect leads to a secondary effect on mitochondrial function. We observed reduced oxygen consumption and ATP production in patient fibroblast cells, indicating compromised oxidative phosphorylation (OXPHOS), underscoring the importance of mtDNA integrity for mitochondrial energy metabolism. Further, we observed reduced MMP and mitochondrial ROS levels, indicating broader disruption of mitochondrial homeostasis. While lower ROS levels in P1 cells may suggest reduced oxidative stress, it is also possible that these cells have an impaired capacity to produce ROS, reflecting more profound dysfunction of the electron transport chain (ETC). We further demonstrate that nucleoid destabilization triggers adaptive remodelling of the mitochondrial network toward hyperfusion, a stress-responsive state that temporarily sustains ATP production and delays mitophagy. However, persistent hyperfusion is associated with impaired mitochondrial dynamics, altered calcium homeostasis, and neurodegenerative features, consistent with the cardiomyopathy and neurological manifestations observed in affected individuals. Collectively, our study establishes the pathogenicity of the MGME1 p. Ala274Thr variant and provides mechanistic insight into how defective processing of replication intermediates drives mtDNA depletion, mitochondrial network remodelling, and multisystem disease. These findings refine the functional role of MGME1 in human mtDNA maintenance and expand the molecular framework underlying MGME1-associated mitochondrial disorders.

## Methods and materials

### Patients

Patients with MGME1 variant were identified from a prospective registry of mitochondrial disorders. They were included in this study after providing written consent. Clinical data were collected using a structured proforma with demographic details, birth and developmental history, antenatal and perinatal risk factors, history of developmental regression, neurological manifestations, consanguinity, and family history. Systemic involvement, including neurological, ophthalmological, cardiac, gastrointestinal, renal, and endocrinological features, was documented. Laboratory evaluations included haematological parameters; liver and renal function tests; blood glucose levels; thyroid function tests; and metabolic investigations, including serum and cerebrospinal fluid lactate, plasma amino acid profile, ammonia, and urine organic acids.

Neurophysiological assessments included electromyography (EMG), electroencephalography (EEG), nerve conduction studies (NCS), visual evoked potentials, and brainstem auditory evoked responses. Magnetic resonance imaging (MRI) of the brain and spine was reviewed to assess stroke-like episodes, Leigh syndrome patterns, and white matter abnormalities, and magnetic resonance spectroscopy (MRS) was performed to assess lactate elevation. Muscle MRI findings were also recorded when performed.

Diagnostic muscle biopsies, when performed, were analysed for the presence of ragged red or ragged blue fibres using modified Gomori trichrome staining and NADH-tetrazolium reductase staining, respectively, succinate dehydrogenase (SDH) staining patterns, and Cytochrome c oxidase (COX), and ultrastructural examination by electron microscopy.

### Next-generation sequencing analysis

Total genomic DNA was extracted from blood samples collected from 5 patients using the PureLink Genomic DNA Mini Kit (Thermo Fisher Scientific, Waltham, USA). The libraries were sequenced to a mean depth of >80–100X on an Illumina sequencing platform. Germline variant identification was performed using the GATK best practice framework via Sentieon(Li & Durbin, 2010). The obtained sequences were aligned to the human reference genome (GRCh38) using BWA, and the resulting alignments were processed with Sentieon for duplicate removal, recalibration, and indel realignment. Variants were called via the Sentieon Haplotype Caller, and the identified germline variants were annotated using the VariMAT pipeline(McLaren *et al*, 2010). Gene annotation was performed using the VEP program against the Ensembl release 104 human gene model(Cunningham *et al*, 2022).

### *In silico* site-directed mutagenesis and stability prediction of MGME1 variant

To study the effects of variation in MGME1, we developed variant structures by substituting alanine with threonine at position 274 (A274T) using an *in-silico* site-directed mutagenesis approach. The apo structure of MGME1 (PDB ID: 5ZYW) was used as a template, and a point mutation (A274T) was introduced via the mutagenesis wizard in PyMOL. Evolutionary conservation of residues was assessed using ConSurf(Ashkenazy *et al*, 2016), which maps sequence conservation onto the protein’s three-dimensional structure. Multiple structure-based protein stability prediction tools, including mCSM(Pires *et al*, 2014a), SDM2.0(Pandurangan *et al*, 2017), DUET(Pires *et al*, 2014b), Dynamut2(Rodrigues *et al*, 2021), PoPMuSiC(Dehouck *et al*, 2011), and HoTMuSiC(Pucci *et al*, 2016), were used to estimate the structural and thermodynamic consequences of the variation. These methods rely on statistical potentials, graph-based signatures, and normal mode analysis to compute the change in folding free energy (ΔΔG) and predict whether a variation is stabilizing or destabilizing.

### All-atom molecular dynamics simulations

To assess the effect of variation in MGME1, all-atom molecular dynamics (MD) simulations were performed using GROMACS v2022.4(Abraham *et al*, 2015) with the CHARMM36 force field(Lee *et al*, 2016). Six systems were developed: wild-type apo MGME1 (PDB: 5ZYW), MGME1 bound to single-stranded DNA (5ZYU), MGME1 bound to 3′ overhang double-stranded DNA (5ZYT), and their respective A274T variant counterparts. Each structure was solvated in an octahedral box of TIP3P water molecules, neutralized with counterions, and adjusted to 0.15 M NaCl. Energy minimization was performed using the steepest descent algorithm for 50,000 steps, followed by equilibration under the NVT and NPT ensembles for 10 ns and 20 ns, respectively. Production runs were carried out for 500 ns with periodic boundary conditions, a 2fs time step, and LINCS constraints on bonds involving hydrogen atoms. The temperature (310 K) and pressure (1 bar) were controlled using a velocity-rescaled thermostat and Parrinello–Rahman barostat, respectively. Post-MD analyses were performed using standard GROMACS tools and in-house scripts to evaluate structural stability, conformational flexibility, and interaction dynamics of the systems.

### Generation of patient-derived skin fibroblast lines

Control, P1, and P2 cells were derived from patients through skin biopsy at Sree Chitra Tirunal Institute for Medical Sciences and Technology, Trivandrum (IEC: 78/2023), and each patient provided written informed consent. Control, P1, and P2 were cultured in advanced DMEM (Gibco, Thermo Fisher Scientific, USA) supplemented with 10% foetal bovine serum (Gibco), and 1% penicillin‒streptomycin (Thermo Fisher Scientific, USA), and 50 μg/ml uridine (Sigma‒Aldrich, Merck) as per previously established protocol(Kandettu *et al*, 2025). When the cells reached 80% confluence, they were trypsinized for subsequent passaging or experimental procedures.

### cDNA conversion and real-time PCR

Total RNA was extracted from Control, P1, and P2 cells using TRIzol Reagent (Thermo Fisher Scientific, USA) according to the manufacturer’s protocol. The concentration and purity of isolated RNA were determined spectrophotometrically by measuring absorbance at 260 nm. RNA was reverse-transcribed to cDNA with the help of the High-Capacity cDNA Reverse Transcription kit (Applied Biosystems, USA) according to the manufacturer’s instructions. Quantitative real-time PCR (qRT-PCR) was performed using PowerUp™ SYBR™ Green Master Mix (Applied Biosystems, USA). The relative expression of *MGME1* was normalized to the internal control, β-Actin. The comparative cycle threshold (2^−ΔΔCt^) was calculated for relative quantification of transcript levels. Data was analyzed using QuantStudio Design and Analysis software (Applied Biosystems, USA) (Supplementary Table 2).

### Immunoblot analysis

Cell lysates from Control, P1, and P2 samples were prepared using RIPA buffer (Sigma-Aldrich, USA) containing a protease inhibitor cocktail (Sigma-Aldrich, USA), and protein concentration was quantified using the Bradford assay (Sigma-Aldrich, USA). Proteins (30–50 µg) were separated on an SDS-PAGE and transferred onto a nitrocellulose membrane. The membrane was then blocked with 5% BSA (Himedia, India) to prevent nonspecific binding and incubated overnight at 4°C with MGME1 polyclonal primary antibodies (1:10000; Proteintech, USA) and the Total OXPHOS Human WB Antibody Cocktail (1:1000; Abcam, UK). After incubation with their respective primary antibodies, the blots were incubated with HRP-conjugated anti-rabbit secondary antibodies (1:10000; Jackson ImmunoResearch Labs, USA) and anti-mouse secondary antibodies (1:10000; Bio-Rad, USA), respectively, for 2h at room temperature. Protein bands were visualized using an enhanced chemiluminescence (ECL) reagent (Bio-Rad Laboratories, USA) on iBright 1500 (Invitrogen, USA). Protein band intensities were analyzed in ImageJ (Fiji) and normalized to β-Actin.

### Immunofluorescence

For immunofluorescence staining, Control and P1 cells were cultured on sterile coverslips at an initial density corresponding to about 60%. Further, the cells were fixed with paraformaldehyde (4%) for 5min and then permeabilized using Triton X-100 (0.5 %) (Sigma-Aldrich, USA). Cells were then blocked with 5 % BSA (Himedia, India) and incubated with the TFAM antibody (1:100; Cell Signalling Technology, USA) overnight at 4 °C. After incubation, cells were then labelled with a FITC secondary antibody (Invitrogen, USA). Cells were stained with Hoechst (Invitrogen, USA) for 5min, washed with PBS, and mounted onto a glass slide using Mowiol (Sigma, USA). The images were captured using an SP8-DMi8 confocal microscope (Leica Microsystems, Germany) at 63× magnification.

### Analysis of mitochondrial DNA copy number

Genomic DNA was extracted from Control, P1, and P2 cells using the PureLink Genomic DNA Mini Kit (Thermo Fisher Scientific, USA). Quantitative real-time PCR (Thermo Fisher Scientific, USA) was performed to analyze mitochondrial DNA content using PowerUp SYBR Green Master Mix (Applied Biosystems, USA). Analysis of mtDNA copy number was determined using mitochondrial gene Cytochrome c oxidase subunit II (*MT-CO2*) and the endogenous control β-Actin.

### Analysis of mitochondrial DNA replication

Control and P1 cells were cultured in a 6-well plate and were exposed to 2mM 2′-3′-dideoxycytidine (ddC, Sigma-Aldrich, USA) for 4 days, after which cells were cultured for an additional 4 days in ddC-free medium. Genomic DNA was collected every alternate day, and quantitative real-time PCR (Thermo Fisher Scientific, USA) was performed on mitochondrial gene Cytochrome c oxidase subunit II (*MT-CO2*) and the endogenous control β-Actin to assess replication efficiency as described previously (Bonora *et al*, 2021).

### Mitochondrial repair assay

Control and P1 cells were grown in six-well plates and treated with 200 µM H₂O₂ (HiMedia, India) for 15 minutes. Following treatment, the cells were either collected immediately or allowed to recover in fresh medium for 6 hours before harvesting. Genomic DNA was then extracted from the cells using the PureLink Genomic DNA Mini Kit (Thermo Fisher Scientific, USA).

For DNA damage analysis, quantitative long-range PCR was performed using the LongAmp Taq PCR Kit (New England Biolabs) with 15 ng of total genomic DNA as the template. The PCR products obtained were then quantified using the PicoGreen double-stranded DNA (dsDNA) assay kit (Thermo Fisher Scientific, USA). Fluorescence intensity was measured using a 200 PRO multimode reader (Tecan, Switzerland). The fluorescence values obtained from the short-fragment PCR products were normalized to those from the long-range PCR products to ensure accurate quantification(Bonora *et al*, 2021).

### Analysis of mitochondrial DNA content, dynamics, and nucleoid structure

For live-cell imaging of mitochondrial DNA, 8×10^4^ Control and P1 cells were seeded on sterile coverslips. After 48 hrs, the cells were fixed with paraformaldehyde (4%) and subsequently stained with 100nM MitoTracker Red CMXRos and Quant-iTPicoGreen dsDNA Reagent (Invitrogen, USA) and incubated for 20 minutes at 37°C in the dark. After incubation, the cells were mounted on glass slides using mowial. Fixed slides prepared from patient-derived fibroblasts were used to analyze mitochondrial length and mitochondrial nucleoids using super-resolution microscopy. Imaging was performed on a Leica Stellaris 8 microscope (Leica Microsystems) equipped with a Leica DMi8 CS scanhead. For STED microscopy, an HC Plan-Apo ×100/1.4 NA STED White oil-immersion objective was used, while point-scanning confocal imaging employed an HC Plan-Apo ×63/1.4 NA oil-immersion objective or an HC Plan-Apo ×63/1.4 NA water-immersion objective for hiPSC-CM imaging. The system was equipped with a pulsed white-light laser (440–790 nm; 78 MHz) with output powers of >1.1 mW at 440 nm, >1.6 mW at 488 nm, >2.0 mW at 560 nm, >2.6 mW at 630 nm, and >3.5 mW at 790 nm, as well as a pulsed 775-nm STED depletion laser. Confocal images were acquired using HyD S or HyD X detectors in either analogue or digital mode, whereas STED imaging was performed exclusively using HyD X detectors in photon-counting mode. Nucleoid numbers were quantified using the Particle Analysis tool in ImageJ FIJI following image binarisation. The total number of nucleoids/cells was automatically calculated within a region of interest (ROI) encompassing the entire mitochondrial network. Analysis was performed on n = 500/cell line across three independent biological replicates. Individual mitochondrial length analysis was performed in Leica LAS X software using the line measurement feature. An average of ∼20-25 mitochondria/cell was measured. Mitochondrial length was analyzed using the line measurement tool in ImageJ after autoscale normalization and was represented on a logarithmic scale. Average mitochondrial length was quantified using ImageJ FIJI. Briefly, the background signal was subtracted, and mitochondrial network images were skeletonized. The analyzed skeleton function in ImageJ FIJI was used to measure mitochondrial length. For each mitochondrion, the sum of all branch lengths was calculated and taken as the total mitochondrial length per image.

### Analysis of the mitochondrial oxygen consumption rate

The total mitochondrial oxygen consumption rate and extracellular acidification rate (OCR and ECAR) were evaluated using a standard protocol from the Seahorse Bioscience XF-24 extracellular flux bioanalyzer (Agilent, USA). Control and P1 cells were harvested, seeded onto XF 8-well microplates, and incubated overnight in a 5% CO_2_ atmosphere at 37°C. Oxygen consumption was subsequently assessed using a Seahorse XF-24 extracellular flux bioanalyzer (Agilent, USA) in conjunction with the Seahorse XF Cell Mito Stress Test(Kandettu *et al*, 2025; Bhattacharjee *et al*, 2026).

### Analysis of mitochondrial membrane potential

To assess mitochondrial membrane potential (MMP), 4 × 10^5^ Control and P1 cells were cultured in 6 cm plates. The cells were trypsinized, washed with PBS, and stained with tetramethylrhodamine methyl ester (TMRM) (Invitrogen, USA) and incubated for 30 minutes at 37°C in the dark. After incubation, fluorescence intensity values were analyzed using fluorescence-activated cell sorting (FACS) (Partec, Germany).

### Analysis of Mitochondrial reactive oxygen species

To analyze mitochondrial reactive oxygen species (ROS), 4 × 10^5^ Control and P1 cells were cultured in a 6 cm plate. The cells were trypsinized, washed with PBS, and stained with MitoSOX Red (Invitrogen, USA). And were incubated for 30 mins at 37°C in the dark. After incubation, fluorescence intensity values were measured using fluorescence-activated cell sorting (FACS) (Partec, Germany).

### Statistical analysis

Statistical analysis was performed using GraphPad Prism. ANOVA and Student’s unpaired t-test. A two-tailed Student’s *t*-test was used to compare differences between two groups. and data were represented as mean ± SD, and a *p*-value less than 0.05 was considered statistically significant.

## Data availability

Data supporting this study’s findings are available from the corresponding author upon reasonable request.

## CRediT authorship contribution statement

**Pranavi Hegde**: Writing – review & editing, Writing – original draft, Visualization, Validation, Methodology, Investigation, Formal analysis, Data curation, Conceptualization. **Amoolya Kandettu**: Visualization, Validation, Methodology, Investigation, Formal analysis, Data curation. **Arpan Bhattacharyya**: Visualization, Validation, Methodology, Formal analysis, Data curation. **Budheswar Dehury**: Software, Formal analysis, Data curation. **Sruthi S Nair**: Patient clinical care and advice. **Rajalakshmi Poyuran**: Clinical diagnosis. **Smily Sharma**: Patient clinical care, diagnosis, and advice. **Benu Brata Das**: Visualization, Validation, Formal analysis. **Soumya Sundaram**: Resources, Formal analysis. **Sanjiban Chakrabarty**: Writing – review & editing, Writing – original draft, Visualization, Validation, Supervision, Resources, Project administration, Methodology, Investigation, Funding acquisition, Formal analysis, Data curation, Conceptualization.

## Acknowledgment

We sincerely acknowledge patients and their families for their consent and participation in this study. We are grateful to all referring physicians who made this work possible. We acknowledge and thank the Department of Health Research (R.11014/33/2023-GIA/HR) and Anusandhan National Research Foundation (ANRF-SUPRA, SPR/2023/000321) for financial support. The infrastructure support and funding from DST-FIST, TIFAC-CORE, DBT-Builder, VGST Karnataka, K-FIST, the Manipal Academy of Higher Education, and the Department of Neurology, Sree Chitra Tirunal Institute for Medical Sciences and Technology, Trivandrum are gratefully acknowledged. We are grateful to the patients for their assistance.

## Ethical approval

This study is approved by the Institutional Ethics Committee of Sree Chitra Tirunal Institute for Medical Sciences and Technology, Trivandrum (IEC:78/2023).

